# Population structure of the endangered Siberian flying squirrel (*Pteromys volans*) revealed by genomic and mitochondrial data

**DOI:** 10.1101/2024.09.13.612918

**Authors:** Fernanda Ito dos Santos, Thomas Lilley, Veronika N. Laine, Stefan Prost, Juri Kurhinen, Hanna Susi, Sergey Gashev, Alexander Saveljev, Svetlana Bondarchuk, Svetlana Babina, Aleksander Shishikin, Dmitry Tirski, Aleksandra Vasina, Vladimir Karpin, Jaana A Kekkonen

## Abstract

The Siberian flying squirrel (*Pteromys volans*) is an arboreal rodent with a distribution range that covers large parts of the Eurasian taiga forest zone. However, extensive forestry has resulted in widespread local population declines and extinctions in recent decades. Even though it is widely distributed in Eurasia, almost nothing is known about its phylogeography. Here we used genome-wide single nucleotide polymorphisms (SNPs) and mitochondrial DNA (mtDNA) to investigate population structure, connectivity and genetic diversity in different sites throughout its distribution. Overall, the species shows low nucleotide diversity and heterozygosity. Locations in Finland, on the western edge of the distribution, had the lowest diversities in genomic SNPs and mtDNA, while individuals in the Far East (Sikhote-Alin, Russia) show the highest diversity. These findings fit with a rapid range expansion from the Far East to the west. We found a strong genetic differentiation between Sikhote-Alin and all other populations investigated, which might warrant a revision of their taxonomic classification. The inferred low genetic diversities at the western edge of their distribution are especially worrisome as they are currently experiencing strong population declines and major habitat changes, which can be especially detrimental when standing variation is low. Thus, stressing the need for the revision of the species conservation status.

## INTRODUCTION

Understanding a species’ genetic diversity in respect to its biogeographical distribution is essential for uncovering patterns of current population structure as well as assessing its potential to adjust to changes in the environment. Combined knowledge on the genetic diversity, ecology, demography and distribution of a species also creates a basis for assessing the need for conservation or management. Large-scale information across the range of a species is needed to help understand local patterns (McGarical et al. 2016, Remm et al. 2017). Connectivity within a landscape enhances dispersal between populations and associated gene flow helps maintain demographic stability, genetic diversity and ultimately their adaptive potential (Cushman et al. 2013, 2016; Diniz et al. 2020). In addition to geographical dispersal barriers, habitat fragmentation can change the structure, composition and function of ecosystems leading to negative impacts on species survival (Fischer and Lindenmayer 2007, Liu et al. 2018, Lino et al. 2019, Püttker et al. 2020). Due to recent human mediated environmental changes, many mammals of the taiga biome suffer habitat losses. Forest-dwelling mammals that have specific habitat requirements and limited dispersal abilities due to unsuitable habitats are considered highly susceptible to habitat loss and fragmentation (Koprowski 2005, McAlpine et al. 2006, Zungu et al. 2020). Moreover, fragmentation and loss of forest habitats may be detrimental especially for arboreal and gliding species that depend on trees for movement as well as a food source and for roosting (Ritchie et al. 2009, van der Ree et al. 2010, Lindenmayer et al. 2011, Makelainen et al. 2016).

The Siberian flying squirrel (*Pteromys volans*) is an arboreal rodent with a distribution range that covers large parts of the Eurasian taiga forest zone. The species primarily inhabits mature spruce-dominated mixed forests, but also younger forests with decaying trees in a standing position to a lesser degree (Hurme et al. 2008, Santangeli et al. 2013) as well as urban forests (Makelainen et al. 2015, 2016). The species’ large distribution range extends from Finland and the Baltic Sea in the west through Siberia to the Pacific coast in the east, also inhabiting the Korean peninsula and Hokkaido, Japan. Over its entire range, the species is listed as Least Concern (LC) with a decreasing population trend on the International Union for Conservation of Nature (IUCN) Red List (Shar et al. 2016). In Russia, the Siberian flying squirrel (listed LC) is on the federal list of game species and hunting is legal in parts of the region (Kurhinen et al. 2011, 2016, Saveljev et al. 2020). However, the species is subject to hunting only formally, since the skins have no utilitarian value. At the same time, the Siberian flying squirrel is in the Red Data Books of 23 (out of 97) regions of Russia (Lissovsky et al. 2019), and even classified as Critically Endangered in some regions. In the east, it is classified as endangered (EN) in South Korea (Lim et al. 2021) and Vulnerable (VU) in China (Jiang et al. 2016). On the western edge of the distribution range their largest population in Finland is listed as Vulnerable (VU) due to vast population declines in the last decades (Hanski 2006, Selonen et al. 2010a), while it is listed as Critically Endangered (CE) in Estonia (Timm and Remm 2011). In Latvia it is considered locally extinct since 2013. A recent study indicated major habitat shrinkage in Belarus (Abramchuk 2021). It might also still be present in the north east of the Ukraine (Abramchuk 2021, Zagorodniuk 2022). The biggest threats for the species are habitat fragmentation, changes in tree compositions, reduction of old-growth forests as well as decreasing amount of decaying wood and loss of suitable habitat due to forestry (Selonen and Makelainen 2017, Jokinen et al. 2019).

The habitat requirements, dispersal and demographics are relatively well-known for the species through extensive ecological studies (Reunanen et al. 2002, Lampila et al. 2009, Selonen et al. 2010b, 2013, Makelainen et al. 2016). Some phylogeographic and genetic diversity studies have also been conducted. Studies on mitochondrial DNA data from individuals across the range indicated division into three genetic lineages (Far Eastern, Hokkaido, and Northern Eurasia) suggesting that glacial refugia of the species would have been associated with forest dynamics in the Pleistocene (Oshida et al. 2005, Lee et al. 2008). Microsatellite data, in turn, has been used to describe genetic differentiation and gene flow between sampling sites in fragmented landscapes at small geographical scales (Nummert et al. 2020). Moreover, in Finland, individuals from regions with greatest declines were found to have the lowest estimates of genetic diversity (Lampila et al. 2009). However, there is still a large gap in knowledge of the current genetic diversity across the entire distribution range and consensus is lacking on what is the taxonomic grouping within the Siberian flying squirrel over its distribution range. Studies based on either morphology or mitochondrial DNA have resulted in estimates between 3 to 10 lineages (Ognev 1940, Oshida et al. 2005, Pavlinov and Lissovsky 2012, Thorington et al. 2012, Gashev et al. 2019). The wide distribution range of the species over the Eurasian continent contains variation in climate, vegetation (Binney et al. 2017) and landscape structure with human alterations (Wallenius et al. 2010). Moreover, there are potential geographical barriers for gene flow such as mountain ranges and rivers (Eidesen et al. 2013, Lee et al. 2015).

Here we used genome-wide single nucleotide polymorphisms (SNPs) data obtained using double-digest restriction site associated DNA sequencing (ddRADseq) to study the genetic diversity and phylogeography of the Siberian flying squirrel using five sampling locations ranging from the western to the eastern edge of its distribution. To understand potential dispersal barriers between populations we use an estimated effective migration model to analyze gene flow among the populations. In addition, we add results from mitochondrial DNA from the same populations to study historical ancestry and species delimitation. As is most often seen among widely distributed species, we predict that genetic differentiation between populations increases with geographical distance (Slatkin 1987). Furthermore, we predict that there are some major dispersal barriers for this species potentially driving its genetic structure. We also expect that populations at the edge of the range have smaller genetic diversity due to founder effect resulting in smaller effective population sizes and greater genetic differentiation from more central populations (Eckert et al. 2008).

## MATERIAL AND METHODS

Samples were obtained from Finland and Russia during the years of 2006 and 2021 and are part of the collection of the Finnish Museum of Natural History (LUOMUS), in Helsinki (Figure 1; Table S1). In Finland, the samples were obtained from several locations and grouped in two subgroups: Western and Eastern Finland (Figure 1; Table S1). We wanted to test whether the eastern and western locations are genetically differentiated as some structuration is suggested for the species in Finland (Lampila et al. 2009), like in other mammal species studied in the country (Tammeleht et al. 2010, Kerminen et al. 2017). In Russia, the Siberian flying squirrel was sampled in three regions: Tyumen, Baikal and Sikhote-Alin (Figure 1; Table S1).

**Figure 1.**
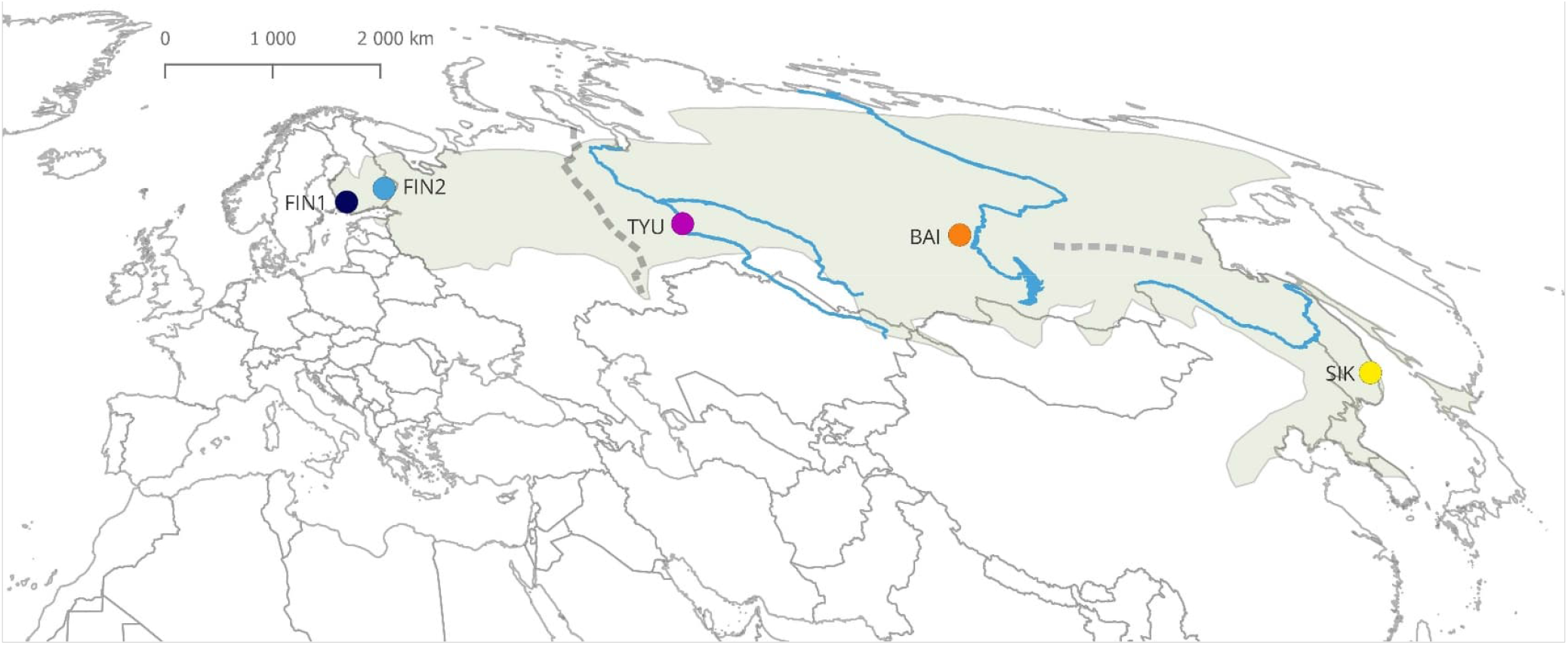
Map of sampling sites and the distribution range of *Pteromys volans* in Eurasia. Living range of the species is presented as a shaded green area. Large rivers Amur, Irtysh, Lena and Ob are presented as blue lines. Ural and Stanovoymountain ranges are represented by gray dashed lines. Sampling sites, represented as colored circles, are as follows: Western Finland (dark blue), Eastern Finland (light blue), Tyumen (purple), Baikal (orange) and Sikhote-Alin (yellow). Map projection: Mollweide Equal Area.

Genomic DNA was extracted from individual frozen tissue samples using the NucleoSpin tissue kit (Macherey-Nagel®) following the manufacturer’s protocol. All DNA stocks were quantified and normalized to 12 ng/µL to be used for molecular processing.

### Genomic data

Double-digestion RADseq libraries were prepared using an adapted protocol for low-concentration samples (Elshire et al. 2011; Lemopoulos et al. 2018), with the restriction enzymes PstI-HF™ and BamHI (New England Biolabs). Sequencing was performed by Bioname Oy (Turku, Finland) using Illumina Novaseq6000 over one lane with 150 paired-end reads. Detailed protocols are available in the Appendix A.

Sequencing quality of the demultiplexed raw reads was assessed using F_AST_QC v.0.11.9 (Andrews 2010). Cleaning and trimming processes were performed using fastp v.0.21.0 (Chen et al. 2018) to remove low-quality bases (< 20 Phred scores) and Illumina adapters. Downstream analyses were performed with Stacks 1.48 (Catchen et al. 2013) and vcftools 4.1 (Danecek et al. 2011). The *de novo* pipeline from Stacks was used for SNP calling, with the default parameters after testing for several combinations of parameter settings (Paris et al. 2017) and assigning the samples to one of the five subpopulations representing the areas they were sampled (Table S1). SNP filtering used the following parameters: maf = 0.01; max-missing = 0.75; minDP = 10 and maxDP = 50.

The final dataset was then used to run *populations* from Stacks to estimate genetic diversity indices for each subpopulation, including nucleotide diversity (π), expected (H_E_) and observed (H_O_) heterozygosity and inbreeding coefficients. To analyze the population structure of the species we used Plink 1.09 (Purcell et al. 2007) to perform a Principal Component Analysis (PCA) and plotted the results for the first three Principal Components (PCs) on R, using the *tidyverse* 2.0 package. The contribution of ancestral populations was estimated with using the mapping files obtained from Stacks, genotype likelihoods obtained with ANGSD 0.940 (Korneliussen et al. 2014), with the options: *- GL 1 -doGlf 2 -doMajorMinor 1 -SNP_pval 1e-6 -doMaf 1*, and ngsAdmix (Skotte et al. 2013). We carried out 20 replicates for each ngsAdmix run ranging from k=2 to k=7 and estimated likelihood scores for the different K.

We used vcftools 4.1 (Danecek et al. 2011) to calculate the pairwise Weir and Cockerham weighted F_ST_ estimates (Weir and Cockerham 1984) for each area sampled. We considered F_ST_ values of 0 to 0.05 to be of low differentiation, and values > 0.15 distinctly differentiated (Hartl and Clark 1997). We also calculated the pairwise geographic distances between the areas with the Haversine method assuming a spherical earth, implemented in the function *distm* in the R package geosphere 1.5.14 (Hijmans et al. 2017), using the latitude and longitude coordinates of the sampling locations. The distances calculated were used to test the isolation by distance (IBD) applying a Mantel test, with 10,000 permutations and considering alfa = 0.05.

We estimated effective migration rates through-out the Siberian flying squirrel’s entire and Finnish distribution, respectively. To do so, we used the mapping files obtained from Stacks and generated genotype likelihoods using ANGSD 0.940 (Korneliussen et al. 2014), with the following parameters: *-GL 1 -doMaf 1 -doMajorMinor 1 - only_proper_pairs 0 -doIBS 1 -doCounts 1 -makeMatrix 1*. These were then used as an input for EEMS (Petkova et al. 2016). We plotted 2,000 demes and carried out 2 million MCMC iterations, with a 10% burn-in. We sampled every 4,999 iterations. The effective migration surface was then plotted using the R package reemsplots2 1.1.0 (https://github.com/dipetkov/reemsplots2) and the CRAN packages rworldmap 1.3-8, rworldxtra 1.01, sf 1.0-15 and ggplot2 3.5.0. Lastly, we inferred a phylogenetic network for all Siberian flying squirrel samples. To do so, we first used ANGSD with the following parameters: -GL 1 -doMaf 2 -doMajorMinor 1 -doGeno 2 -doPost 1 -doSaf 1 -fold 1 - SNP_pval 1e-6, to generate genotypes and genotype probabilities. These were then converted into the adegenet input format to calculate genetic distances using the R package adegenet (Jombart and Ahmed 2011) using PopGenTools (https://github.com/CGRL-QB3-UCBerkeley/PopGenTools). The phylogenetic network was then visualized using Splitstree (Huson et al. 2008).

### Mitochondrial data

Two mitochondrial DNA markers were used to analyze the species genetic diversity and structure: a fragment of 420 bp of the cytochrome B (CytB) gene; and 561 bp of the mitochondrial control region sequence (D-loop). The PCR reactions had total volume of 10 µL, containing 5 µL of Phusion high-fidelity PCR master mix (Thermo Fisher Scientific), 1 µM of each primer, 1 µL of extracted DNA (25-50 ng/µL) and 2 µL of mQ water. Following the program: activation step at 98°C for 10 s; 39 cycles of 98°C for 1 s, 55°C for 5 s and 72°C for 15 s; and a final extension step at 72°C for 2 min. Primers for both markers were selected based on Nummert et al. (2020): 5’-CGA TTC TTC GCA TTC CAT TT-3’ and 5’-TAG TTG GCC GAT GAT GAT GA-3’ for CytB; and 5’-TGC ACA GCC CCA TTA ATA CA-3’ and 5’-GGG AGG GTT TCG AGT CAA AT-3’ for D-loop. Positive reactions were purified and diluted 1:3, and then sequenced with ABI 3730 Genetic analyzer (Applied Biosystems, Foster City, CA). Sanger sequencing was performed in Molecular Ecology and Systematics laboratory (University of Helsinki).

The sequences were edited and aligned with Geneious Prime 2023.2.1 (www.geneious.com). Genetic diversity indexes were estimated for each subpopulation using DnaSP v.6 (Rozas et al. 2017) and included: haplotype diversity (h), nucleotide diversity (π) and Tajima’s D test (Tajima 1989). Pairwise genetic distances between the subpopulations were calculated with MEGA11 (Tamura et al. 2021) using both datasets separately. To visualize relationships between individuals from all sampled subpopulations, a median-joining haplotype network was constructed in PopArt 1.7 (Leigh and Bryant 2015) using a concatenated dataset for both markers. To investigate the subdivision of genetic variation within and between sampling locations, analysis of molecular variance (AMOVA) was performed with ARLEQUIN 3.5.2.2. (Excoffier and Lischer 2010). Furthermore, discriminant analysis of principal components (DAPC; Jombart et al. (2010)) using the freely available R package ‘adegenet’ (Jombart 2008). R version 4.3.0 (R Core team 2013) was used for the analysis. The *read*.*dna* function of the ‘ape’ package was used to import our mtDNA sequence data into R. The function *DNA-bin2genind* (‘adegenet’) was used to convert the data into a genind class object. This genind object was then used as the input for the DAPC analysis using the *dapc* function. Twelve principal components were retained and 2 discriminant analysis eigenvalues. The 2D plot was generated using the *scatter* function (‘adegenet’). To run the species delimitation analyses, first we estimated a Bayesian phylogenetic reconstruction. To do so, we concatenated sequences of CytB and D-loop and determined the best substitution model with jModelTest 2.1.6 (Darriba et al., 2012). We then performed three independent runs with BEAST v.1.10.4 (Suchard et al. 2018), for 10 million generations, sampling every 1,000 and with a 10% burn-in. The convergence of the runs was checked with TRACER 1.5 (Drummond and Rambaut, 2007), and LogCombiner 2.4.5 and TreeAnnotator 2.4.5 (Drummond and Rambaut, 2007) were used to combine the runs and generate the best tree. The tree was then used to perform two tree-based coalescent species delimitation approaches: Generalized Mixed Yule Coalescent model (GMYC; Fujisawa and Barraclough 2013) and Bayesian Poisson Tree Processes model (bPTP; Zhang et al. 2013). We used the R package *splits* 1.0.19 (Ezard et al. 2009) to run the single-threshold GYMC analysis, and the bPTP web server to run the highest Bayesian supported solution with default parameters settings (Zhang et al. 2013). The Automatic Barcode Gap Discovery (ABGD; Puillandre et al. 2012) method does not require a phylogenetic reconstruction and was also performed with the default settings and Kimura K2p nucleotide substitution model at the web server (Puillandre et al. 2012).

## RESULTS

In total, 76 individuals of Siberian flying squirrel were sampled from five different locations (Figure 1; Table S1).

### SNP data

After sequence read quality filtering, mapping, SNP calling and filtering, 28,218 biallelic genotypes were retained for all 76 individuals and used in downstream analyses. In general, nucleotide diversity is very low in all populations sampled, ranging from 0.00482 to 0.19316 (Table 1). The populations in Finland had the lowest and the easternmost population (SIK) the highest values with regards to nucleotide diversity (Table 1). The heterozygosity is also low for all populations analyzed, and follows the same pattern as the nucleotide diversity, decreasing from East to West across the sampled area (Table 1; Figure 1). However, the species also show low inbreeding coefficients, with negative values (Table 1).

**Table 1.** Genetic diversity of Siberian flying squirrels sampled in Finland and Russia based on SNP data, including number of individuals (N), nucleotide diversity (π), inbreeding coefficient, and the expected (HE) and observed (HO) heterozygosity.

| Location | N | Nucleotide diversity | Inbreeding coefficient | Heterozygosity |  |
| --- | --- | --- | --- | --- | --- |
|  |  |  |  | Observed | Expected |
| **Western Finland (FIN1)** | **15** | 0.00509 | -0.00400 | 0.00736<br>($\pm 0.00517$) | 0.00491<br>($\pm 0.00155$) |
| **Eastern Finland (FIN2)** | **21** | 0.00482 | -0.00391 | 0.00710<br>($\pm 0.00507$) | 0.00470<br>($\pm 0.00149$) |
| **Tyumen (TYU)** | **13** | 0.01289 | -0.00266 | 0.01463<br>($\pm 0.00667$) | 0.01225<br>($\pm 0.00321$) |
| **Baikal (BAI)** | **14** | 0.04390 | 0.00360 | 0.04349<br>($\pm 0.01591$) | 0.04222<br>($\pm 0.01242$) |
| **Sikhote-Alin (SIK)** | **13** | 0.19316 | 0.06616 | 0.17095<br>($\pm 0.03096$) | 0.18495<br>($\pm 0.02718$) |

In the PCA, the first two principal components explain 51.33% of the total variation and show a clear pattern of substructure within our sampling (Figure 2). The most separated group is formed by the individuals from Sikhote-Alin, another group is formed by the samples from Baikal, and the third group is formed by the samples from Tyumen and Finland (Figure 2). This pattern can also be observed for the PCs 2 and 3, which together explain 15.55% of the total variation (Figure S1). However, when only the individuals from Finland are considered, the PCA shows no sign of structuring (Figure S2).

**Figure 2.**
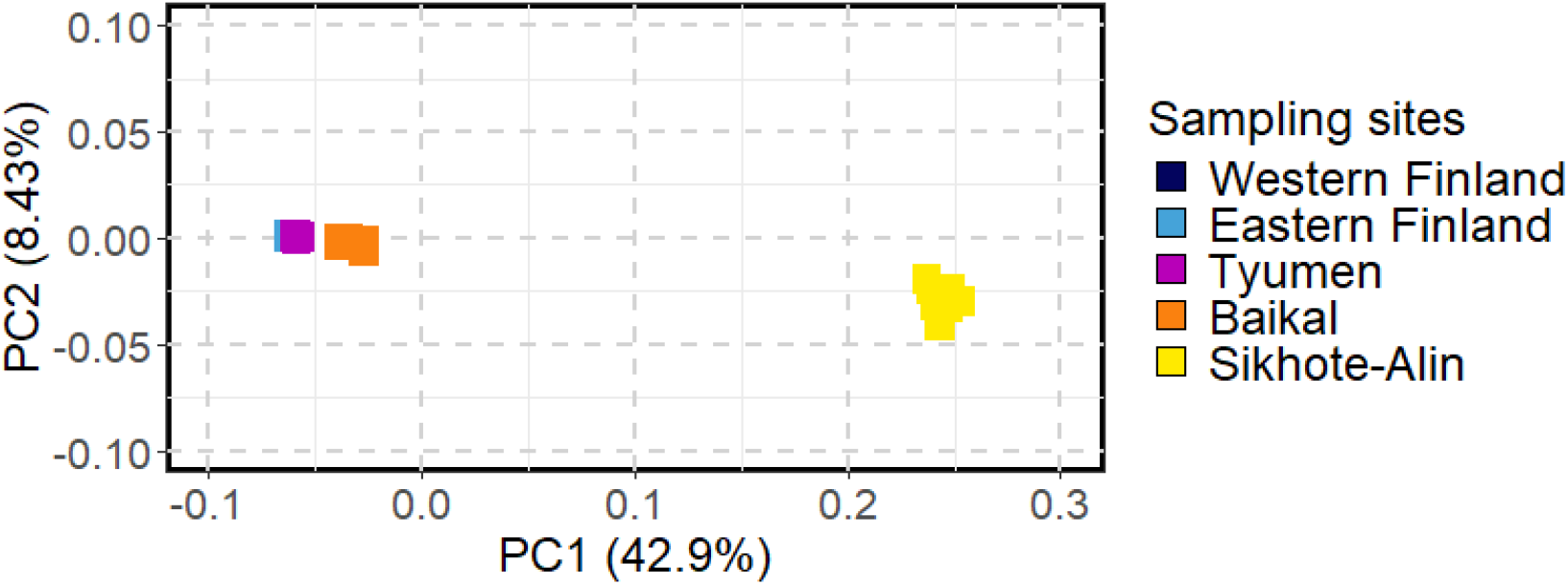
Genetic variation of *Pteromys volans* from Finland and Russia explained by the first two principal components and based on 28,218 SNPs for 76 individuals.

Admixture analyses with two ancestral groups (K=2) separates Sikhote-Alin from all other populations (Figure 3) and presents Baikal sharing some ancestry with Sikhote-Alin. At three ancestral groups (K=3), Sikhote-Alin, Baikal and all the others form separate groups, with Baikal showing shared ancestry with the group comprising Finland and Tyumen populations. At K=4, Tyumen separates from Finland and at K=5 no additional geographic splitting occurs. Both in K=4 and K=5, Baikal shows shared ancestry with Tyumen. Based on likelihood scores calculated for the different number of ancestral groups, K = 4 shows the lowest variance (Figure 4). Overall, K=2-5 show very little variance.

**Figure 3.**
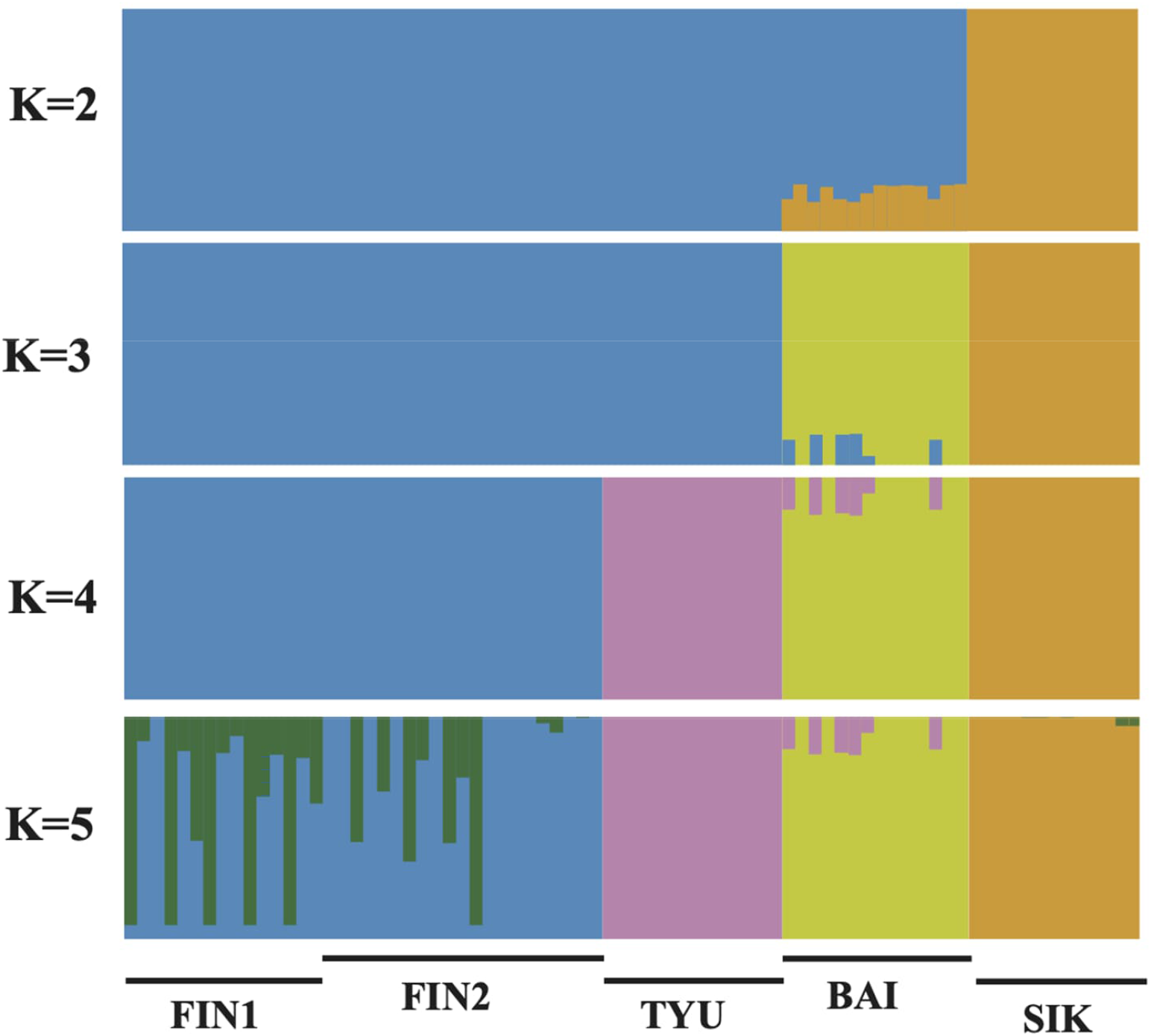
Ancestral populations (K=2 to K=5) of *Pteromys volans* from Finland and Russia estimated with ngsAdmix using the SNP data. The number of ancestral populations is represented by K. Individuals are represented by the vertical bars and grouped according to the sampling area. Colors do not represent the same population for each value of K.

**Figure 4.**
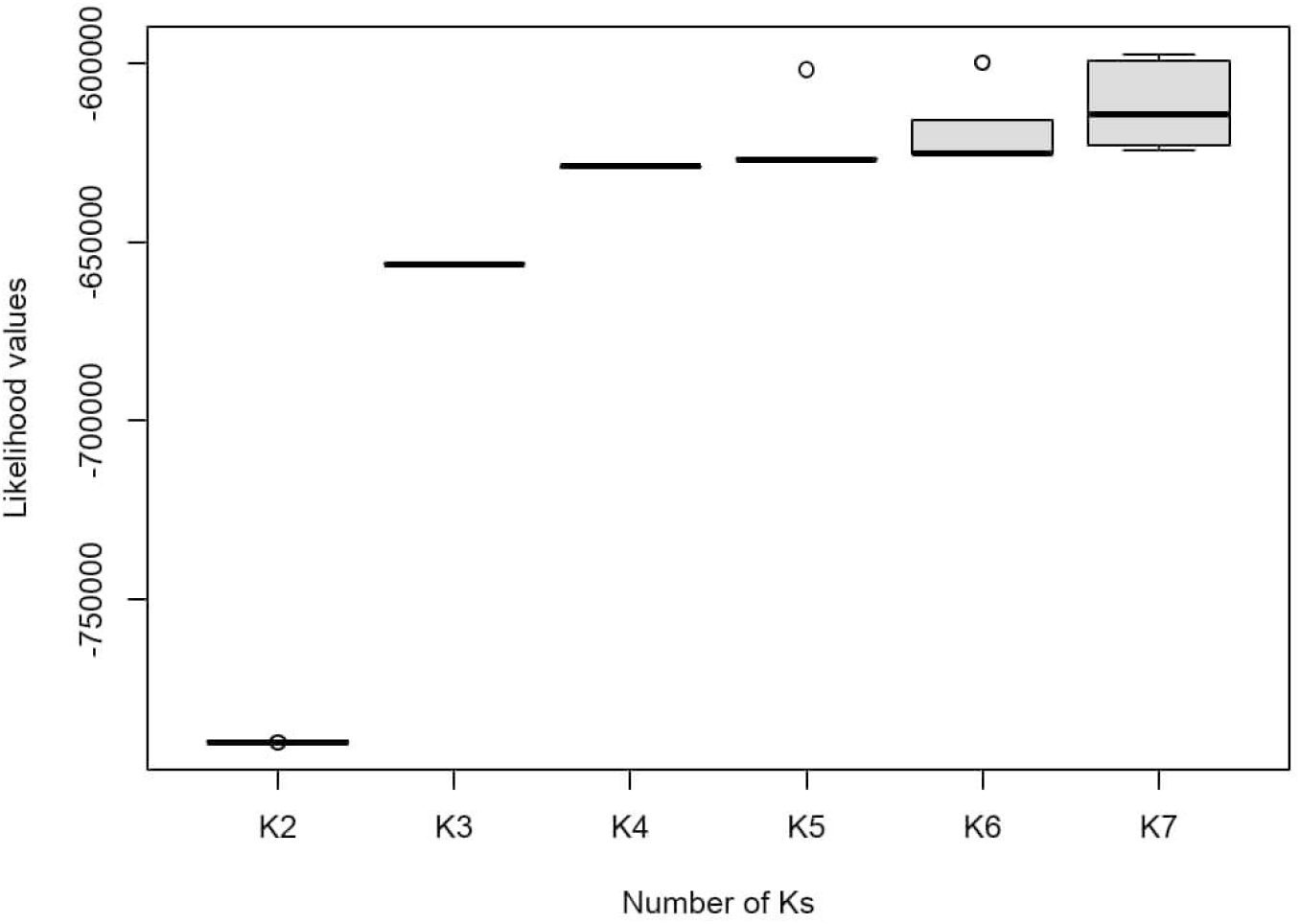
Likelihood scores for estimating the best K-value of admixture model. Outliers represented by circles.

Overall pairwise F_ST_ values are very high, most of them above 0.40 (Table 2). Only the two sampling locations in Finland present low genetic differentiation, with F_ST_ values lower than 0.05. Eastern Finland and Sikhote-Alin present the biggest genetic differences, with F_ST_ value of 0.610 (Table 2). The pairwise geographical distances range from 363 km between the locations in Finland to 6757 km between the westernmost location in Finland (FIN1) and the easternmost location in Russia (SIK; Table 2). Mantel test suggests that the genetic and geographical distances are strongly correlated (r = 0.7736; p = 0.02). Therefore, the greater the geographical distance, the higher the genetic differentiation.

**Table 2.** Pairwise Weir and Cockerham weighted F_ST_ estimate (above diagonal) based on SNP data and geographic distance in kilometers (below diagonal) for the five sampling locations of Siberian flying squirrels from Finland and Russia.

|  | **FIN1** | **FIN2** | **TYU** | **BAI** | **SIK** |
| --- | --- | --- | --- | --- | --- |
| **FIN1** | - | 0.024 | 0.479 | 0.496 | 0.562 |
| **FIN2** | 363 | - | 0.499 | 0.539 | 0.610 |
| **TYU** | 2514 | 2183 | - | 0.373 | 0.470 |
| **BAI** | 4463 | 4124 | 2237 | - | 0.445 |
| **SIK** | 6757 | 6425 | 4614 | 2388 | - |

The EEMS results support our findings, showing that gene flow is facilitated among both Finnish sampling locations (FIN1 and FIN2) and Tyumen, while an increased resistance to gene flow can be observed among Tyumen and Baikal, and between Baikal and Sikhote-Alin (Figure 5). In the phylogenetic network all locations form separate groups, except the two Finnish ones which make a single group (Figure 6).

**Figure 5.**
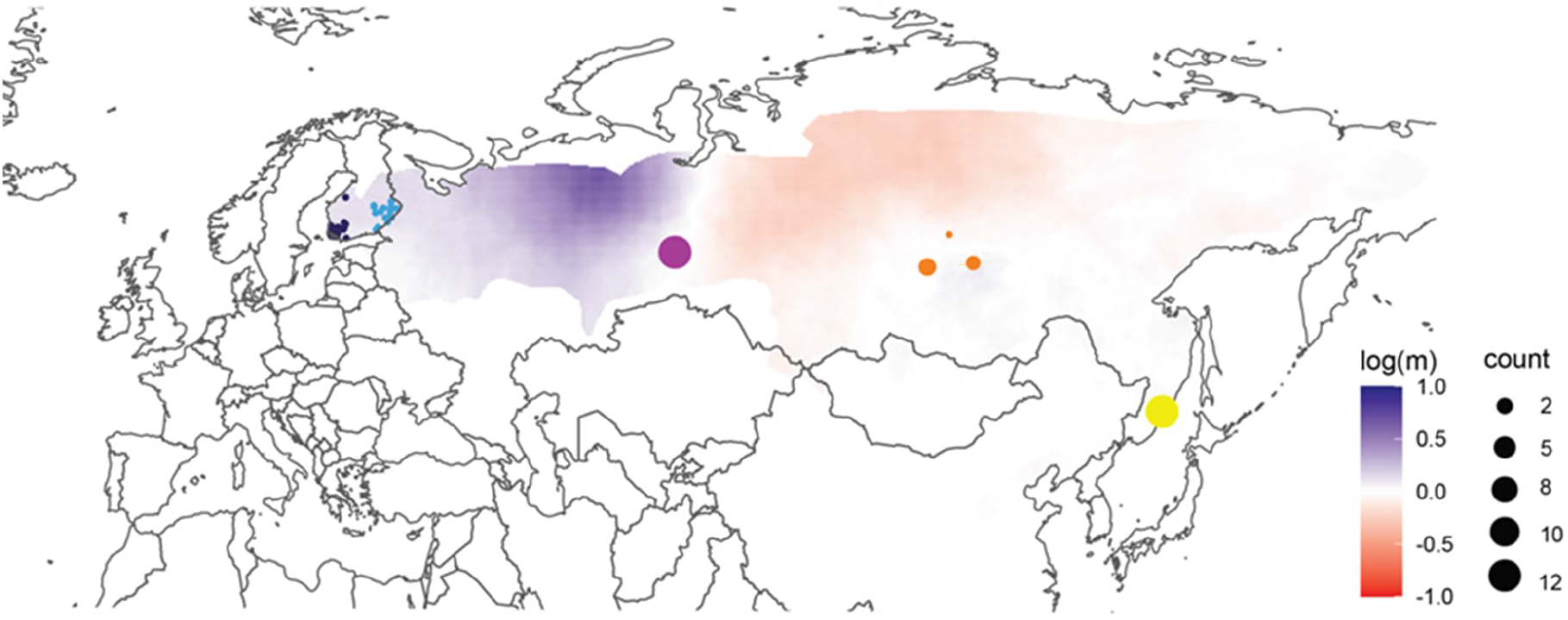
Effective migration surfaces estimated using EEMS (Petkova et al. 2106). Circles indicate sample locations (scaled by the number of individuals): Western Finland (dark blue), Eastern Finland (light blue), Tyumen (purple), Baikal (orange) and Sikhote-Alin (yellow); colors reflect effective migration rates relative to the overall migration rate found throughout the distribution of Siberian flying squirrels.

**Figure 6.**
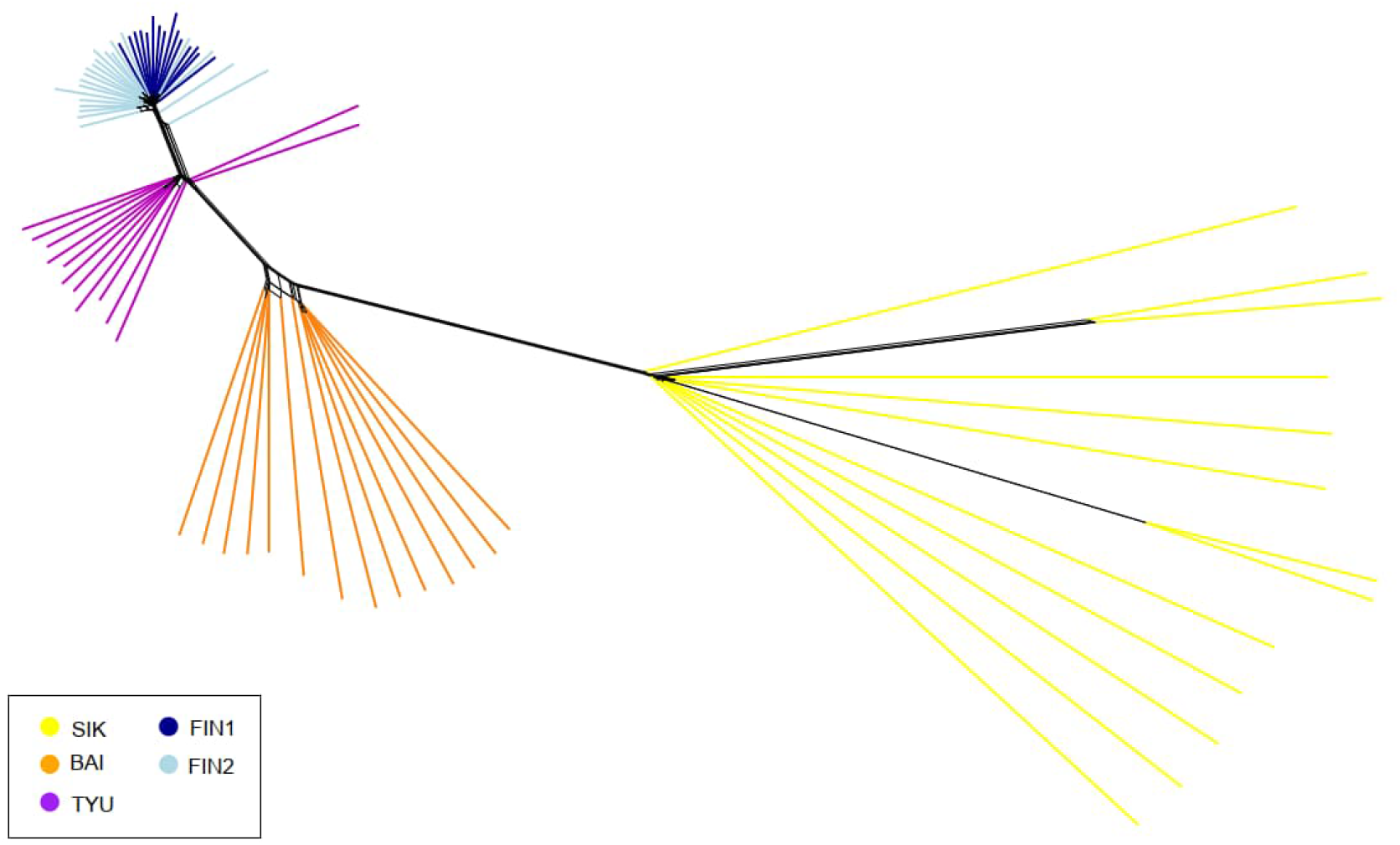
Phylogenetic network of the Siberian flying squirrels based on 28,218 SNPs. Individual samples are colored according to the sampling area.

### Mitochondrial data

Sequences of CytB (420 bp) and D-loop (561 bp) were generated for all 76 individuals. In general, the analysis showed very low genetic diversity in the analyzed area (Table 3). For CytB, the haplotype diversities ranged from 0 to 0.923 and nucleotide diversities from 0 to 0.0074, with Western Finland (FIN1) presenting the lowest values. For D-loop haplotype diversities ranged from 0.593 to1 and nucleotide diversities from 0.0014 to 0.0183. Tajima’s D statistic was negative for all locations but failed to reject the null hypothesis of neutrality as they were all non-significant. For the concatenated data the diversity indices are very similar to the ones for the individual genes (Supplementary Table S2).

**Table 3.** Diversity indices of *Pteromys volans* based on mitochondrial data (CytB and D-loop). The number of haplotypes (N), haplotype diversity *(h)*, nucleotide diversity *(π)* and Tajima’s D neutrality test are presented for each sampling location. None of the Tajima’s D neutrality test p-values were significant.

| Location | CytB |  |  |  | D-loop |  |  |  |
| --- | --- | --- | --- | --- | --- | --- | --- | --- |
| | N | ($h$) | ($\pi$) | Tajima's D | N | ($h$) | ($\pi$) | Tajima's D |
| **Western Finland (FIN1)** | 1 | 0.000<br>($\pm 0.000$) | 0.0000<br>($\pm 0.000$) | 0.000 | 5 | 0.593<br>($\pm 0.021$) | 0.0014<br>($\pm 0.000$) | -0.565 |
| **Eastern Finland (FIN2)** | 2 | 0.091<br>($\pm 0.007$) | 0.0002<br>($\pm 0.000$) | -1.162 | 6 | 0.649<br>($\pm 0.008$) | 0.0020<br>($\pm 0.000$) | -1.573 |
| **Tyumen (TYU)** | 3 | 0.410<br>($\pm 0.024$) | 0.0010<br>($\pm 0.000$) | -0.909 | 10 | 0.962<br>($\pm 0.002$) | 0.0048<br>($\pm 0.000$) | -0.317 |
| **Baikal (BAI)** | 10 | 0.923<br>($\pm 0.004$) | 0.0074<br>($\pm 0.000$) | -1.070 | 14 | 1.000<br>($\pm 0.001$) | 0.0170<br>($\pm 0.000$) | -0.475 |
| **Sikhote-Alin (SIK)** | 5 | 0.705<br>($\pm 0.015$) | 0.0049<br>($\pm 0.000$) | -0.344 | 12 | 0.987<br>($\pm 0.001$) | 0.0183<br>($\pm 0.000$) | -0.624 |

The haplotype network was constructed based on the concatenated mitochondrial data (981 bp) and shows the same pattern as previous analyses (Figure 7). Samples from Sikhote-Alin form a distinct group that does not share haplotypes with any of the other groups. Baikal, despite its geographical proximity with Sikhote-Alin, shows haplotypes closely related with Finland and Tyumen, but also presents exclusive haplotypes. On the other hand, all Finnish samples were grouped together, with the individuals from Tyumen being closely related to them (Figure 7).

**Figure 7.**
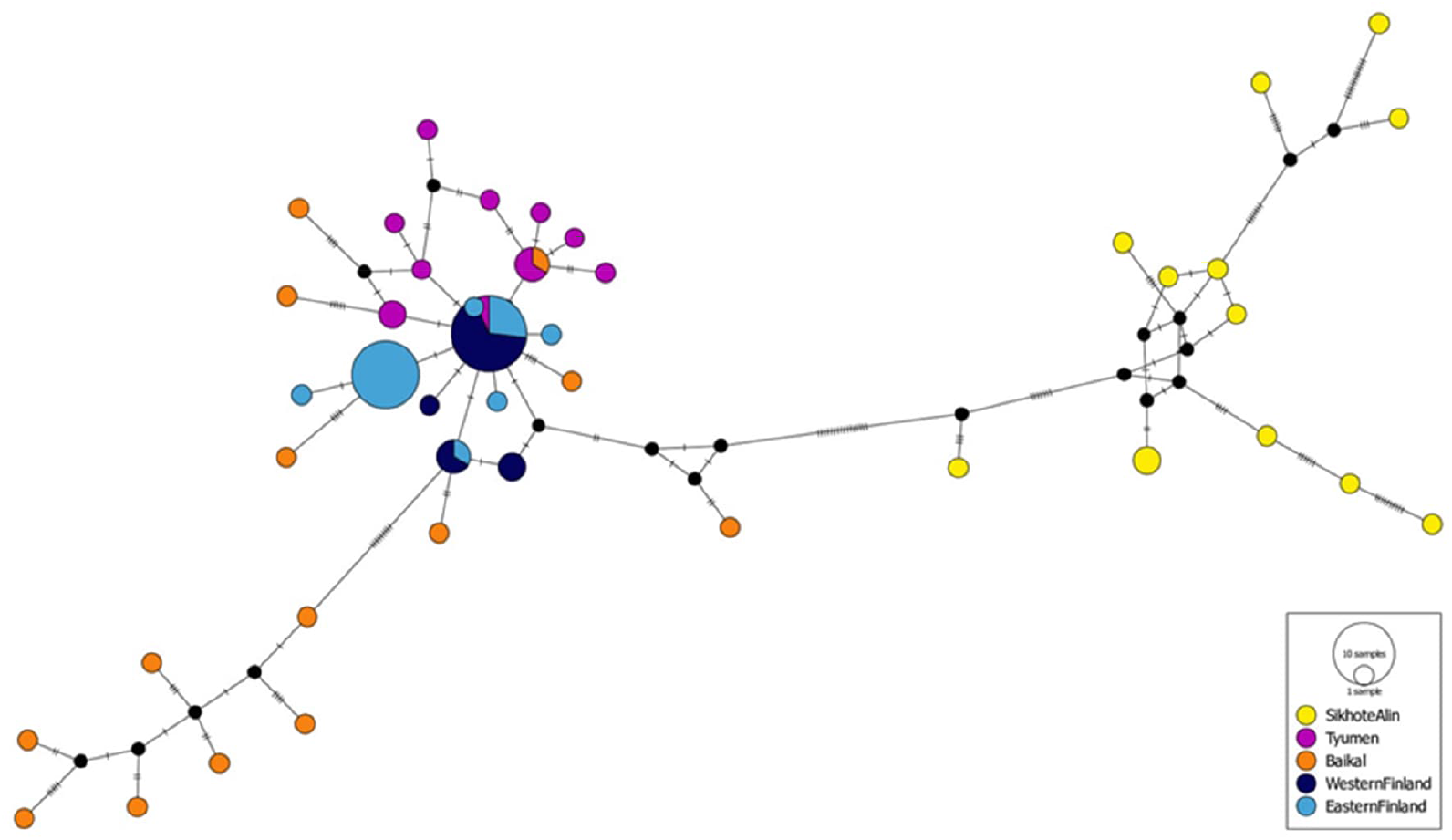
Median-joining network presenting haplotype relationships of the mtDNA dataset (CytB and D-loop). Haplotypes are colored according to their geographical distribution. Circle sizes are proportional to the haplotype frequency. Hatch marks represent the number of mutations between nodes. Missing haplotypes are indicated by black circles.

Genetic distances show that all other populations were furthest apart from Sikhote-Alin, even Baikal which is geographically closest. Genetic distances for CytB ranged from 0 to 0.016 and for D-loop from 0.002 to 0.043 (Table 4). Analysis of molecular variance of the concatenated mtDNA sequences shows that most of the variation occurs between groups of locations when Sikhote-Alin is considered separate from the rest of the sampling locations (72,92%) (Table 5). AMOVA for CytB and D-loop separately show similar results (Supplementary table S3).

**Table 4.** Genetic distance matrices of mitochondrial sequences of CytB (below diagonal) and D-loop (above diagonal) for 76 individuals of *Pteromys volans* from Finland and Russia.

|  | FIN1 | FIN2 | TYU | BAI | SIK |
| --- | --- | --- | --- | --- | --- |
| **FIN1** | - | 0.002 | 0.004 | 0.013 | 0.039 |
| **FIN2** | 0.000 | - | 0.005 | 0.014 | 0.041 |
| **TYU** | 0.001 | 0.001 | - | 0.015 | 0.042 |
| **BAI** | 0.005 | 0.005 | 0.005 | - | 0.043 |
| **SIK** | 0.016 | 0.016 | 0.016 | 0.016 | - |

**Table 5.** Analysis of molecular variance for the five subpopulations of *Pteromys volans*, based on concatenated mtDNA sequences (CytB and D-loop). AMOVA ran for all locations separately and grouping Sikhote-Alin against all the other locations.

| Source of variation | All locations separately |  |  |  |  |
| --- | --- | --- | --- | --- | --- |
|  | df | SS | Variance components | % of variation | *p* |
| Among locations | 4 | 267.38 | 4.26 | 61.19 | 0.000 |
| Within locations | 71 | 191.91 | 2.70 | 38.81 |  |

| Sikhote-Alin vs rest of the locations |  |  |  |  |  |
| --- | --- | --- | --- | --- | --- |
| Among groups | 1 | 219.97 | 9.55 | 72.92 | 0.194 |
| Among locations within groups | 3 | 47.41 | 0.84 | 6.44 | 0.000 |
| Within locations | 71 | 191.91 | 2.70 | 20.64 | 0.000 |

The discriminant analysis of principal components (DAPC) of the concatenated mtDNA sequences showed the same clustering as the PCA analysis of the ddRAD data (see Figure 2) with a close relationship of Finland and Tyumen, the next closest population being Baikal, while Sikhote-Alin showed high differentiation to all other populations (Figure 8). The DAPC for CytB and Dloop separately gave similar results (Supplementary Figure S3).

**Figure 8.**
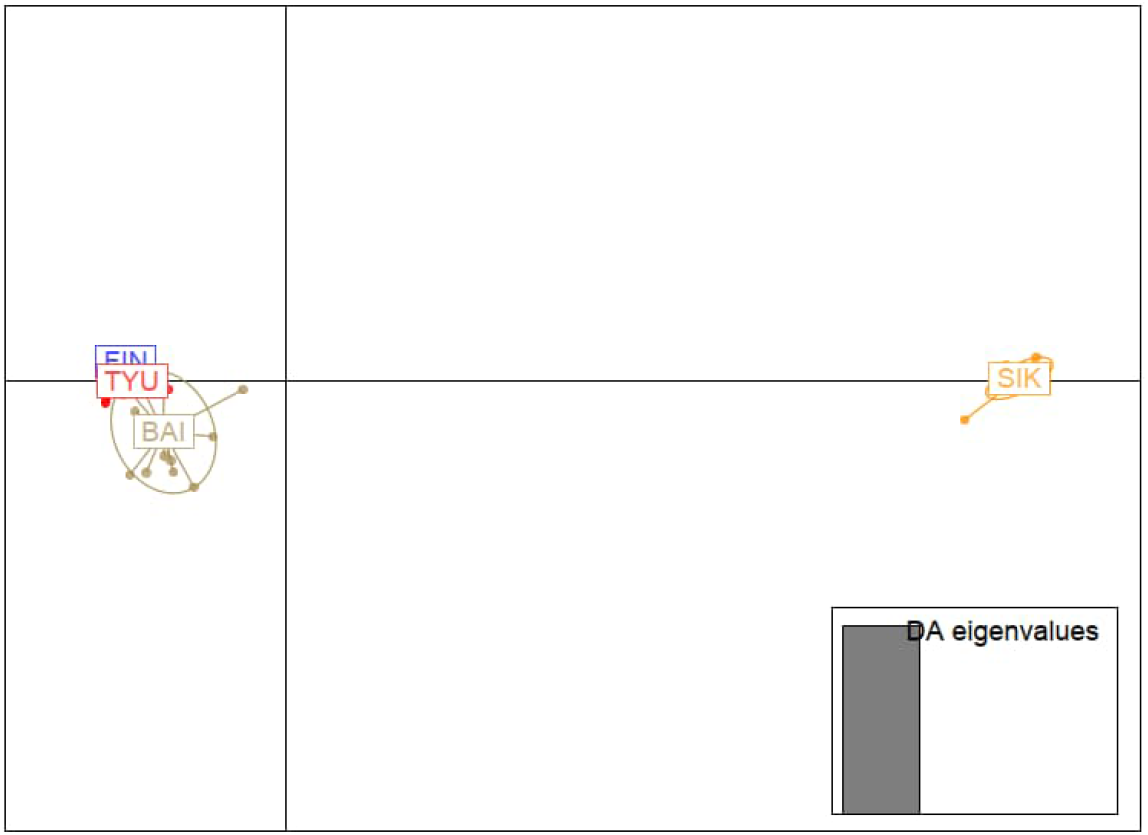
DAPC clustering of the concatenated mitochondrial sequences (CytB and Dloop). The plot shows a strong differentiation of SIK from all other populations. It further shows a closer relationship of FIN to TYU than to BAI. Twelve principal components were retained and 2 discriminant analysis eigenvalues.

The results of three species delimitation analyses indicate the existence of distinct lineages in the Siberian flying squirrel within the sampled area (Supplementary Figure S4). All methods indicated an exclusive lineage for Sikhote-Alin, that could represent a putative subspecies. A second potential sublineage comprises a few individuals from the Baikal area. Although the methods suggest at least one more lineage comprising all the samples from Finland and Tyumen, and a few samples from Baikal, the support for these clades is very weak and the methods also disagree on these relationships.

## DISCUSSION

Here, we present new information on the genetic diversity, connectivity and phylogenetic distribution of the Siberian flying squirrel. Overall, we found low nucleotide diversity and heterozygosity in all locations except Sikhote-Alin (both in mitochondrial and nuclear DNA). Interestingly, there was a clear pattern of genetic diversity being lowest in Finland and increasing towards the East, with the highest values in Sikhote-Alin. The expected heterozygosities and nucleotide diversities were 2 to 38 times higher in Tyumen, Baikal and Sikhote-Alin than in Finland (Sikhote-Alin having around 4 times higher values than any other location). This trend fits a rapid range expansion from the East to the west where populations at the leading edge of the range expansion exhibit lowest diversities (Excoffier et al. 2009, Mona et al. 2014). In addition, high values in Sikhote-Alin may be explained by a low human impact on the forests of the sampling location as well as a large nature reserve area close by (Kurhinen, unpubl).

The levels of genetic diversity in mtDNA were very similar to the nuclear SNPs: Finland had the lowest haplotype and nucleotide diversities and Sikhote-Alin the highest. The mtDNA results were coherent with previous studies on Siberian flying squirrels. Both Oshida et al. (2005) and Lee et al. (2008) found similar patterns of the northwest Eurasian group having clearly lower mtDNA diversity compared to the Far Eastern group. The low diversity in Finland found in this study is also of the same magnitude as in Nummert et al. (2020), where low nucleotide and haplotype diversities in CytB and D-loop in a small Estonian squirrel population were presented. The study also included populations from Finland with very similar results as we obtained in the present study: haplotype diversity for CytB 0.125 and nucleotide diversity 0.00027 and for D-loop 0.642 and 0.00156. In our results, haplotype diversity for CytB was 0.056 and nucleotide diversity 0.00013 and for D-loop 0.724 and 0.0021.

In addition to differences in genetic diversities over the range, we discovered a high genetic distance between the sampling locations and pronounced isolation-by-distance. F_ST_ values between population pairs for SNPs were on average 0.45 and the Mantel test showed strong correlation between genetic and geographical distances. However, despite the large spatial distances and levels of genetic differentiation, in several phylogenetic analyses Finland and Tyumen grouped surprisingly close together as well as with Baikal whereas Sikhote-Alin differentiated clearly. This is interesting also in terms of the relatively low dispersal ability of the species (maximum recorded natal dispersal distance of only 9km; Hanski and Selonen 2009) and requirements for arboreal habitat. Most of the analyses grouped Tyumen and Finland close to each other, indicating that Finnish individuals share ancestry and/or connectivity with Tyumen. In the phylogenetic network based on SNPs, the Finnish group was closest to Tyumen while separating more from Baikal and Sikhote-Alin. The admixture analysis found similar divisions into ancestral groups as the other analysis. The likelihood scores for admixture analysis give high support for all K=3-5 divisions, K=4 showing the lowest variance separating Sikhote-Alin, Baikal, Tyumen and Finland combined. In the AMOVA for mtDNA, the largest part of variation was between groups where Sikhote-Alin was analyzed against the rest of the locations. Moreover, in DAPC all other locations grouped together while there was no overlap between them and Sikhote-Alin. EEMS analysis supported these results as it showed gene flow between Finnish locations and Tyumen, while an increasing resistance to gene flow was observed between Tyumen and Baikal. Based on these results, the Ural mountains do not seem to create a considerable dispersal barrier between Tyumen and Finland. The Ural mountains have a lot of forested areas thus providing suitable habitats for the Siberian flying squirrel. The river Ob may present some dispersal barrier between Tyumen and Baikal. Wide rivers can act as geographical barriers as the gliding distance of the species is limited. Average gliding distances of 10 to 20m and up to 50m have been reported for this species (Asari, Yanagawa and Oshida 2007). Furthermore, river banks may not have suitable trees and man-made bridges do not offer gliding possibilities. In addition, the world’s largest peatland, Vasyugan swamp, lies between Tyumen and Baikal potentially acting as a barrier. In our analyses, Sikhote-Alin was clearly separated from all the other locations. When considering the Sikhote-Alin population, in addition to the effects from post-glacial expansion, some biogeographical barriers may contribute to the strong genetic differentiation. Firstly, Sikhote-Alin has a different climate (“monsoon”) and a conifer-broadleaved forest type compared to the continental climate of other sampling locations and taiga forest type. Secondly, the 700 km Stanovoy Mountain range lies northeast of Lake Baikal passing into the Dzhugjur ridge in the east. A significant part of the territory of the Stanovoy Range and the adjacent mountain areas has vast treeless areas (mountain tundra), dwarf cedar forests and glaciers, which are more difficult for the Siberian flying squirrel to overcome than the forested Ural Mountains. Thus, the Stanovoy Range, together with Lake Baikal and large Lena and Amur rivers, likely represents a geographic barrier to the Siberian flying squirrel. Results from this study are similar to those of Oshida et al. 2005 and Lee et al. 2008, who found large-scale patterns separating lineages for the Far East and northern Eurasia as well as Hokkaido. They speculated that during the last glaciation multiple refugia could have been formed in Eurasia leading to these genetic lineages.

Both similar and differing patterns of genetic diversity and structuring have been found in other studies on taiga rodents expanding to northern Eurasia after the last glacial period. Fedorov et al. 2008 studied the mtDNA phylogeography of the wood lemming *(Myopus schisticolor)* which inhabits boreal forest of the Eurasian taiga zone with a relatively similar range as the Siberian flying squirrel. The study found otherwise low genetic structure between the study locations but one major phylogeographic discontinuity in southeastern Siberia. They also compared phylogeographic structures across a taxonomically diverse array of other taiga species revealing similar patterns. In five out of six species a major discontinuity was found in southeastern Siberia and the Far East implicating two different refugial areas. The harvest mouse *(Micromys minutus)* also appears to have a similar pattern (Pilevich et al. 2023) as found in our study despite not being solely a forest species. The harvest mice lineage in the Far East is clearly separated from the rest of Russia which in turn was different from European lineages but closer to the European lineage than the Far East lineage. For a more comprehensive knowledge on the current genetic structure of Siberian flying squirrels more sampling locations covering the areas between the locations used here would be needed. In addition, to establish the genetic lineages it would be beneficial to combine morphological information with genetic data. For instance, Ognev (1940) and Gashev et al. (2019) found fur color (whiteness and shade of red color) to be an indicator of subclade.

As the Siberian flying squirrel is declining in many parts of the range, conservation actions based on comprehensive information are needed. Results from this study combined with previous ones may warrant a taxonomic review. Sackett et al. (2014) introduced five criteria for evaluating putative subspecies which could be taken into more detailed consideration also for Siberian flying squirrels. Results from our study indicate that the first criteria of *genotypic separation of putative subspecies* is met as both PCA and DAPC distinguished Sikhote-Alin as separate from the rest of the locations. The second criteria of *spatial segregation* is difficult to assess as continental Siberian flying squirrels have an overall continuous distribution and more detailed studies are needed to assess barriers of gene-flow between the existing populations. The third criteria calls for *monophyly in the phylogenetic tree*. This pattern is observed in the haplotype network of mtDNA data and the phylogenetic network based on SNP data. The fourth criteria of *comparing differentiations* we did not test. The fifth criteria is based on whether a relatively *large fraction of genetic variation is partitioned between putative subspecies*. Our AMOVA results are congruent with this. These criteria have been used for recommendations in e.g. Gunnison’s prairie dog and cheetah conservation regarding translocations (Sackett et al. 2014, Prost et al. 2022). For Siberian flying squirrels, these types of issues may become urgent in the western edge of its distribution where we found the genetic diversity to be extremely low.

### Siberian flying squirrel conservation in Finland and the European Union

Our findings highlight the importance of understanding patterns of genetic diversity in Finland in relation to the patterns over the range. Due to low genetic diversity, their ability to adapt to changes on a genetic level is also likely lowered thus making them susceptible to adverse effects of habitat change. Therefore, because Finland has the largest Siberian flying squirrel population within the European Union, it has a particular responsibility for the conservation of the species. Vast declines of the species in the European side of its range have led it to be listed in Annexes II and IV of the EU Habitat Directive and is thus seen as a priority species within the European Union. In Finland, the species has declined drastically in the last years (locally 20-58% declines within 10-20 years) resulting in the classification as Vulnerable (VU) on the Red List (Hyvärinen et al. 2019). Estonia’s small population is listed as Critically Endangered (CR). The forest habitat requirements (preferring of mature spruce-dominated mixed forests, standing decaying trees) of the species inevitably create conflicts with forestry interests and thus the Siberian flying squirrel has been included into political discussion in Finland (Selonen and Makelainen 2017). The species has become a flagship species for taiga forests of Eurasia (Selonen and Makelainen 2017) potentially increasing interest in conservation of the species itself as well as conservation of taiga biome in general. Moreover, as the Siberian flying squirrel has rather specific habitat requirements for mature boreal forests it has been suggested to have umbrella and indicator species characteristics (Hurme et al. 2008) meaning that conserving it may potentially conserve other species with similar habitat requirements.

## ACKNOWLEDGEMENTS

We thank all the collectors for the samples provided. We thank the Finnish Functional Genomics Centre supported by University of Turku, Åbo Akademi University and Biocenter Finland for allowing the sequencing of our samples. JAK thanks the MES lab personnel for the support.

## DATA ACCESSIBILITY

All reads and sequences are available on NCBI under Bioproject PRJNA1183735. Complete scripts used for the analyses can be accessed at github.

