## Supplementary material for "Population structure of the endangered Siberian flying squirrel (*Pteromys volans*) revealed by genomic and mitochondrial data" (from another version of this preprint: version 1, 2024-09-19)

**Table S1.** Sample information of *Pteromys volans* sampled in Finland and Russia. Museum IDs correspond to the Luomus IDs from the Finnish Museum of Natural History (LUOMUS) and can be accessed at laji.fi with all the corresponding data for each sample.

| Sample | Museum ID | Country | Locality | Latitude | Longitude | Subpopulation | Sex |
|---|---|---|---|---|---|---|---|
| FIN1_5984 | http://id.luomus.fi/HT.27330 | Finland | Nousiainen | 60.6125 | 22.1386 | Western Finland (FIN1) | Male |
| FIN1_5983 | http://id.luomus.fi/KS.90908 | Finland | Humppila | 60.9601 | 23.3231 | Western Finland (FIN1) | Male |
| FIN1_5982 | http://id.luomus.fi/HT.27295 | Finland | Huittinen | 61.0857 | 22.6744 | Western Finland (FIN1) | Male |
| FIN1_5980 | http://id.luomus.fi/HT.28925 | Finland | Raasepori | 60.1582 | 23.887 | Western Finland (FIN1) | Male |

| | | | | | | | |
|---|---|---|---|---|---|---|---|
| FIN1_5978 | http://id.luomus.fi/KS.28885 | Finland | Evijärvi | 63.622778 | 23.81 | Western Finland (FIN1) | Male |
| FIN1_5976 | http://id.luomus.fi/HT.27329 | Finland | Nousiainen | 60.6125 | 22.1386 | Western Finland (FIN1) | Female |
| FIN1_5975 | http://id.luomus.fi/HT.27317 | Finland | Lempäälä | 61.3428 | 23.7973 | Western Finland (FIN1) | Female |
| FIN1_5964 | http://id.luomus.fi/KS.28884 | Finland | Eurajoki | 61.2047 | 21.7404 | Western Finland (FIN1) | Male |
| FIN1_5959 | http://id.luomus.fi/KS.28882 | Finland | Lieto | 60.61813 | 22.47784 | Western Finland (FIN1) | Male |
| FIN1_5958 | http://id.luomus.fi/HT.27316 | Finland | Masku | 60.5297 | 22.1222 | Western Finland (FIN1) | Male |

| | | | | | | | |
|---|---|---|---|---|---|---|---|
| FIN1_5955 | http://id.luomus.fi/HT.27322 | Finland | Tampere | 61.4938 | 23.5883 | Western Finland (FIN1) | Male |
| FIN1_5952 | http://id.luomus.fi/HT.26901 | Finland | Urjala | 61.0505 | 23.6194 | Western Finland (FIN1) | Male |
| FIN1_5951 | http://id.luomus.fi/HT.26783 | Finland | Parikkala | 61.5085 | 29.6057 | Western Finland (FIN1) | Male |
| FIN1_5950 | http://id.luomus.fi/KS.90907 | Finland | Loimaa | 60.9805 | 23.0276 | Western Finland (FIN1) | Female |
| FIN1_5949 | http://id.luomus.fi/KS.90905 | Finland | Jokioinen | 60.8964 | 23.4958 | Western Finland (FIN1) | Female |
| FIN2_5990 | http://id.luomus.fi/HT.26780 | Finland | Outokumpu | 62.7258 | 29.0125 | Eastern Finland (FIN2) | Female |

| | | | | | | | |
|---|---|---|---|---|---|---|---|
| FIN2_5989 | http://id.luomus.fi/HT.26804 | Finland | Kontiolahti | 62.6484 | 30.1272 | Eastern Finland (FIN2) | Female |
| FIN2_5988 | http://id.luomus.fi/KS.82993 | Finland | Kuopio | 62.9346 | 27.6607 | Eastern Finland (FIN2) | Female |
| FIN2_5987 | http://id.luomus.fi/KS.82994 | Finland | Kuopio | 62.8365 | 27.698 | Eastern Finland (FIN2) | Female |
| FIN2_5986 | http://id.luomus.fi/HT.26808 | Finland | Heinävesi | 62.4239 | 28.6331 | Eastern Finland (FIN2) | Female |
| FIN2_5985 | http://id.luomus.fi/HT.26781 | Finland | Savonlinna | 61.9125 | 29.2792 | Eastern Finland (FIN2) | Female |
| FIN2_5981 | http://id.luomus.fi/HT.26806 | Finland | Outokumpu | 62.7253 | 29.0446 | Eastern Finland (FIN2) | Female |

| | | | | | | | |
|---|---|---|---|---|---|---|---|
| FIN2_5979 | http://id.luomus.fi/HT.26799 | Finland | Kitee | 62.0284 | 29.858 | Eastern Finland (FIN2) | Female |
| FIN2_5977 | http://id.luomus.fi/HT.26784 | Finland | Kontiolahti | 62.7001 | 30.1032 | Eastern Finland (FIN2) | Male |
| FIN2_5974 | http://id.luomus.fi/HT.27314 | Finland | Lappeenranta | 60.862 | 27.9808 | Eastern Finland (FIN2) | Male |
| FIN2_5973 | http://id.luomus.fi/HT.26800 | Finland | Joensuu | 62.7963 | 30.1541 | Eastern Finland (FIN2) | Female |
| FIN2_5970 | http://id.luomus.fi/HT.26797 | Finland | Joensuu | 62.4334 | 29.9703 | Eastern Finland (FIN2) | Female |
| FIN2_5968 | http://id.luomus.fi/HT.26789 | Finland | Kontiolahti | 62.7635 | 30.0004 | Eastern Finland (FIN2) | Female |

| | | | | | | | |
|---|---|---|---|---|---|---|---|
| FIN2_5967 | http://id.luomus.fi/HT.26786 | Finland | Kitee | 61.9211 | 30.1379 | Eastern Finland (FIN2) | Male |
| FIN2_5963 | http://id.luomus.fi/HT.26875 | Finland | Varkaus | 62.3157 | 27.8947 | Eastern Finland (FIN2) | Female |
| FIN2_5961 | http://id.luomus.fi/HT.26858 | Finland | Lieksa | 63.2569 | 30.0743 | Eastern Finland (FIN2) | Female |
| FIN2_5960 | http://id.luomus.fi/HT.34591 | Finland | Lappeenranta | 61.0727 | 28.2527 | Eastern Finland (FIN2) | Female |
| FIN2_5957 | http://id.luomus.fi/HT.26788 | Finland | Joensuu | 62.4447 | 29.9665 | Eastern Finland (FIN2) | Female |
| FIN2_5956 | http://id.luomus.fi/HT.26787 | Finland | Lieksa | 63.3372 | 29.9408 | Eastern Finland (FIN2) | Female |

| | | | | | | | |
|---|---|---|---|---|---|---|---|
| FIN2_5954 | http://id.luomus.fi/HT.27297 | Finland | Kuopio | 62.8357 | 27.6992 | Eastern Finland (FIN2) | Female |
| FIN2_5953 | http://id.luomus.fi/HT.26805 | Finland | Ilomantsi | 62.6713 | 30.9323 | Eastern Finland (FIN2) | Female |
| TYU_1 | http://id.luomus.fi/LR.2828 | Russia | Tyumen | 58.963889 | 69.260833 | Tyumen (TYU) | Male |
| TYU_3-2 | http://id.luomus.fi/LR.2829 | Russia | Tyumen | 58.963889 | 69.260833 | Tyumen (TYU) | Male |
| TYU_4 | http://id.luomus.fi/LR.2830 | Russia | Tyumen | 58.963889 | 69.260833 | Tyumen (TYU) | Male |
| TYU_6 | http://id.luomus.fi/LR.2831 | Russia | Tyumen | 58.963889 | 69.260833 | Tyumen (TYU) | Female |
| TYU_7 | http://id.luomus.fi/LR.2832 | Russia | Tyumen | 58.963889 | 69.260833 | Tyumen (TYU) | Male |

| | | | | | | | |
|---|---|---|---|---|---|---|---|
| TYU_8 | http://id.luomus.fi/LR.2833 | Russia | Tyumen | 58.963889 | 69.260833 | Tyumen (TYU) | Female |
| TYU_9 | http://id.luomus.fi/LR.2834 | Russia | Tyumen | 58.963889 | 69.260833 | Tyumen (TYU) | Female |
| TYU_10 | http://id.luomus.fi/LR.2835 | Russia | Tyumen | 58.963889 | 69.260833 | Tyumen (TYU) | Male |
| TYU_11 | http://id.luomus.fi/LR.2836 | Russia | Tyumen | 58.963889 | 69.260833 | Tyumen (TYU) | Male |
| TYU_12 | http://id.luomus.fi/LR.2837 | Russia | Tyumen | 58.963889 | 69.260833 | Tyumen (TYU) | Female |
| TYU_16 | http://id.luomus.fi/LR.2838 | Russia | Tyumen | 58.963889 | 69.260833 | Tyumen (TYU) | Female |
| TYU_18 | http://id.luomus.fi/LR.2839 | Russia | Tyumen | 58.963889 | 69.260833 | Tyumen (TYU) | Female |
| TYU_20 | http://id.luomus.fi/LR.2840 | Russia | Tyumen | 58.963889 | 69.260833 | Tyumen (TYU) | Male |

| | | | | | | | |
|---|---|---|---|---|---|---|---|
| BAI_1 | http://id.luomus.fi/LR.2841 | Russia | Baikal | 54.874444 | 105.1 | Baikal (BAI) | Female |
| BAI_2 | http://id.luomus.fi/LR.2842 | Russia | Baikal | 60.453611 | 107.141111 | Baikal (BAI) | Female |
| BAI_3 | http://id.luomus.fi/LR.2843 | Russia | Baikal | 57.766667 | 104.174167 | Baikal (BAI) | Female |
| BAI_7 | http://id.luomus.fi/LR.2844 | Russia | Baikal | 58.066944 | 110.443611 | Baikal (BAI) | Male |
| BAI_10 | http://id.luomus.fi/LR.2845 | Russia | Baikal | 57.766667 | 104.174167 | Baikal (BAI) | Male |
| BAI_12 | http://id.luomus.fi/LR.2846 | Russia | Baikal | 58.066944 | 110.443611 | Baikal (BAI) | Female |
| BAI_15 | http://id.luomus.fi/LR.2847 | Russia | Baikal | 58.066944 | 110.443611 | Baikal (BAI) | Female |
| BAI_18 | http://id.luomus.fi/LR.2848 | Russia | Baikal | 57.766667 | 104.174167 | Baikal (BAI) | Female |

| | | | | | | | |
|---|---|---|---|---|---|---|---|
| BAI_21 | http://id.luomus.fi/LR.2849 | Russia | Baikal | 58.066944 | 110.443611 | Baikal (BAI) | Male |
| BAI_22 | http://id.luomus.fi/LR.2850 | Russia | Baikal | 58.066944 | 110.443611 | Baikal (BAI) | Female |
| BAI_24 | http://id.luomus.fi/LR.2851 | Russia | Baikal | 58.066944 | 110.443611 | Baikal (BAI) | Female |
| BAI_26 | http://id.luomus.fi/LR.2852 | Russia | Baikal | 57.766667 | 104.174167 | Baikal (BAI) | Female |
| BAI_30 | http://id.luomus.fi/LR.2853 | Russia | Baikal | 58.066944 | 110.443611 | Baikal (BAI) | Female |
| BAI_34 | http://id.luomus.fi/LR.2854 | Russia | Baikal | 58.066944 | 110.443611 | Baikal (BAI) | Female |
| SIK_1 | http://id.luomus.fi/LR.2815 | Russia | Sikhote-Alin | 45.476546 | 136.559758 | Sikhote-Alin (SIK) | Male |

| | | | | | | | |
|---|---|---|---|---|---|---|---|
| SIK_2 | http://id.luomus.fi/LR.2816 | Russia | Sikhote-Alin | 45.476546 | 136.559758 | Sikhote-Alin (SIK) | Male |
| SIK_3 | http://id.luomus.fi/LR.2817 | Russia | Sikhote-Alin | 45.476546 | 136.559758 | Sikhote-Alin (SIK) | Female |
| SIK_4 | http://id.luomus.fi/LR.2818 | Russia | Sikhote-Alin | 45.476546 | 136.559758 | Sikhote-Alin (SIK) | Male |
| SIK_5 | http://id.luomus.fi/LR.2819 | Russia | Sikhote-Alin | 45.476546 | 136.559758 | Sikhote-Alin (SIK) | Male |
| SIK_6 | http://id.luomus.fi/LR.2820 | Russia | Sikhote-Alin | 45.476546 | 136.559758 | Sikhote-Alin (SIK) | Female |
| SIK_7 | http://id.luomus.fi/LR.2821 | Russia | Sikhote-Alin | 45.476546 | 136.559758 | Sikhote-Alin (SIK) | Female |

| | | | | | | | |
|---|---|---|---|---|---|---|---|
| SIK_8 | http://id.luomus.fi/LR.2822 | Russia | Sikhote-Alin | 45.476546 | 136.559758 | Sikhote-Alin (SIK) | Female |
| SIK_9 | http://id.luomus.fi/LR.2823 | Russia | Sikhote-Alin | 45.476546 | 136.559758 | Sikhote-Alin (SIK) | Female |
| SIK_10 | http://id.luomus.fi/ LR.2824 | Russia | Sikhote-Alin | 45.476546 | 136.559758 | Sikhote-Alin (SIK) | Male |
| SIK_11 | http://id.luomus.fi/LR.2825 | Russia | Sikhote-Alin | 45.476546 | 136.559758 | Sikhote-Alin (SIK) | Female |
| SIK_12 | http://id.luomus.fi/LR.2826 | Russia | Sikhote-Alin | 45.476546 | 136.559758 | Sikhote-Alin (SIK) | Male |
| SIK_13 | http://id.luomus.fi/LR.2827 | Russia | Sikhote-Alin | 45.476546 | 136.559758 | Sikhote-Alin (SIK) | Female |

**Table S2.** Diversity indices of concatenated mitochondrial sequence of CytB and D-loop. Number of haplotypes, haplotype diversity *(h)*, nucleotide diversity *(π)* and Tajima's D neutrality test are presented for each subpopulation. None of the Tajima's D neutrality test p-values were significant.

| Subpopulation | Number of haplotypes | Haplotype diversity *(h)* | Nucleotide diversity *(π)* | Tajima's D |
|---|---|---|---|---|
| **Western Finland (FIN1)** | 4 | 0,552 (±0,137) | 0,00082 (±0,00024) | -0,395 |
| **Eastern Finland (FIN2)** | 7 | 0,657 (±0,104) | 0,00128 (±0,00039) | -1,688 |
| **Tyumen (TYU)** | 10 | 0,970 (±0,044) | 0,00292 (±0,00047) | -0,890 |
| **Baikal (BAI)** | 14 | 1,000 (±0,027) | 0,01276 (±0,00095) | -0,670 |
| **Sikhote-Alin (SIK)** | 12 | 0,987 (±0,035) | 0,01253 (±0,00157) | -0,593 |

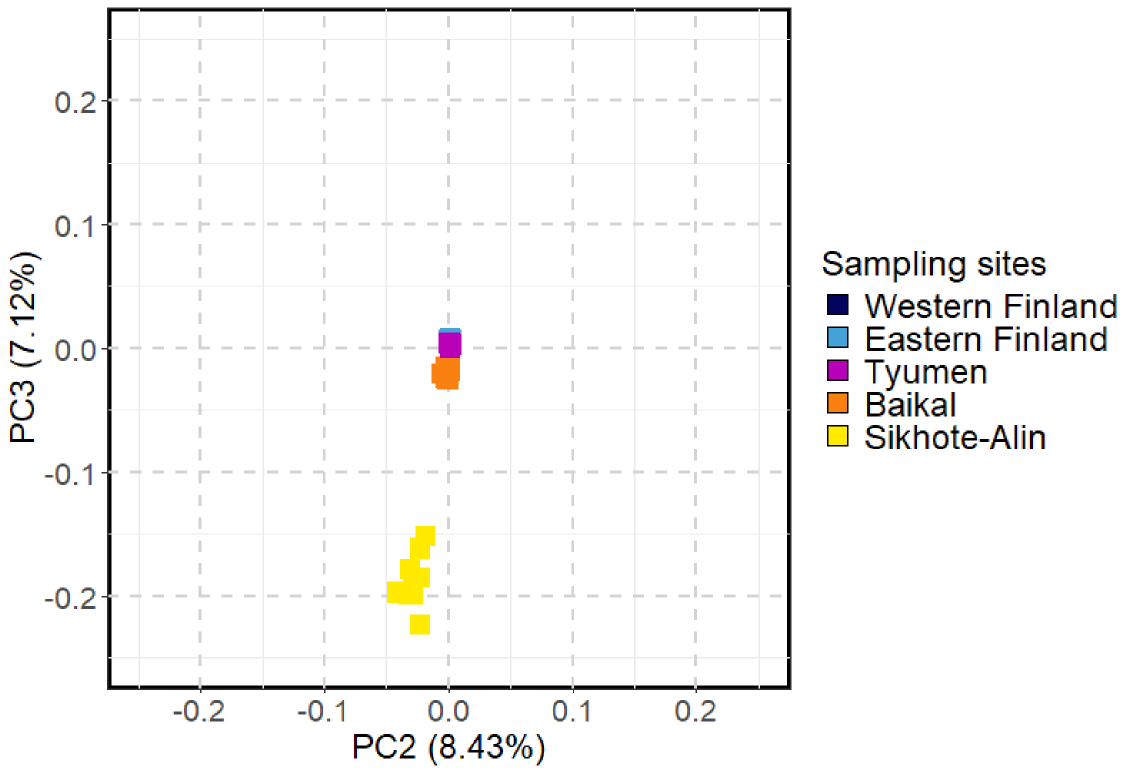


**Figure S1.** Principal components analysis results for *Pteromys volans* from Finland and Russia. Genetic variation from SNP data obtained from 76 individuals explained by the PCs 2 and 3.

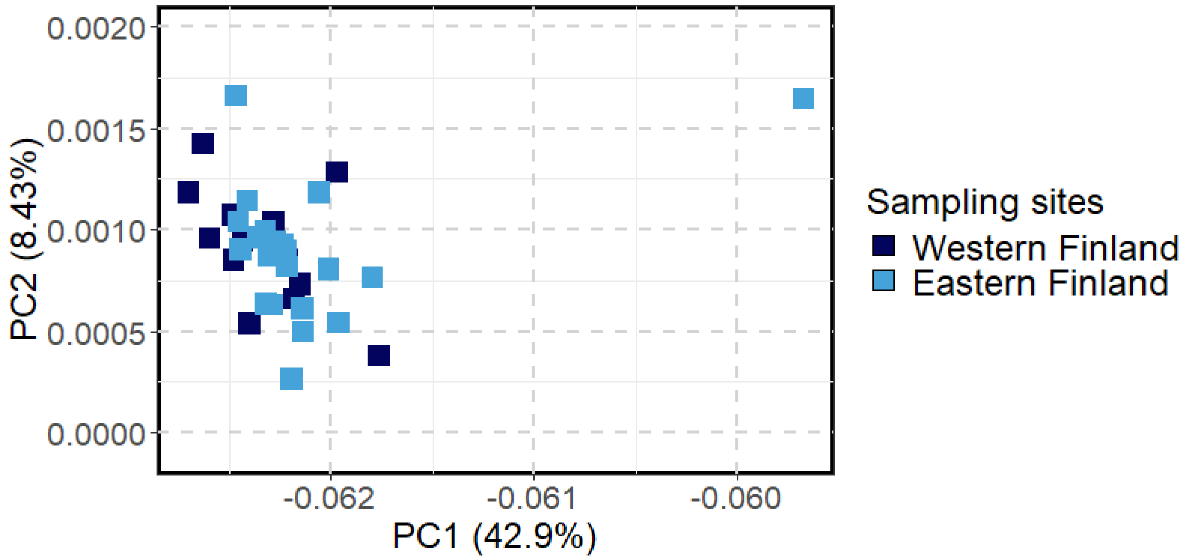


**Figure S2.** Genetic variation of *Pteromys volans* from Finland explained by SNP data from 36 individuals.

**REPORT ddRADseq**

1. SAMPLE PREPARATION

• Altogether 263 DNA samples in microcentrifuge tubes or plates were delivered to Bioname.

• Original DNA concentrations were provided by the customer.

2. DOUBLE RESTRICTION ENZYME DIGESTION

• DNA libraries were prepared by using library preparation protocol adapted from Lemopoulos et al (2018) and Elshire et al (2011) with some modifications.

• In the digestion step, 180 ng of each DNA extract was used in a reaction together with 20 U (units) of each of the restriction enzymes PstI-HF™ and BamHI (New England Biolabs, Catalog numbers: PstI-HF® R3140S; BamHI R0136S) and 2 µl of CutSmart Buffer (provided with the restriction enzymes).

• Digestion was initiated by incubating the mixture for 2 h at 37 °C and terminated by heat-inactivation for 15 min at 75 °C.

3. LIGATION

• The ligation was set up using custom-tailored indexed adapters and a ligase enzyme. The forward (left) primers included a sample-specific unique indices and were compatible with PstIHI™. The reverse (right) primers were compatible with BamHI did not include indices.

• The adapter sequences were for forward (left) adapter top (P1.1): 5′-ACACTCTTTCCCTACACGACGCTCTTCCGATCTxxxxTGCA-3’ and bottom (P1.2) 5′- yyyyAGATCGGAAGAGCGTCGTGTAGGGAAAGAGTGT-3’, and for reverse (right) adapter top (P2.1): 5′-GATCAGATCGGAAGAGCGGTTCAGCAGGAATGCCGAG-3’ and bottom (P2.2): 5′-CTCGGCATTCCTGCTGAACCGCTCTTCCGATCT3’ where “xxxx” denote a 4-8 bp barcode and “yyyy” barcode complement sequences. Altogether, we used 96 different indices. In the ligation step the total reaction volume was 30 µl, consisting of 12 µl digested DNA, 720 U of T4 ligase (New England Biolabs, Catalog number M0202; 400,000 units/ml), 3 µl 10x T4 DNA ligation buffer (provided with the ligase enzyme), 0.5 pmol of both forward and reverse indexed adapters and 11.2 µl of RNase-free H2O.

• Ligation was initiated by incubating the mixture for 1 h at 22 °C and terminated by heat-inactivation for 30 min at 65 °C.

4. NGS LIBRARY PREPARATION

• In the pooling step, each set of 96 uniquely indexed samples were pooled by combining 10 µl of each sample.

• Each sample pool was purified using four columns of Qiagen QIAquick PCR purification kit (Qiagen Corporation, Catalog number 28104), eluting each column in 40 µl.

• To confirm the purification success, the DNA concentration of each pool was measured by Qubit Fluorometer 2.0 device using dsDNA HS assay kit (ThermoFisher Scientific).

• The pools were then PCR-amplified using tagged primers for adding Illumina platform -specific adapters. Primer sequence for forward and reverse (lowercase parts attach to the Illumina flow cell):

5′-aatgatacggcgaccaccgagatctacacTCTTTCCCTACACGACGCTCTTCCGATCT-3’ 5′-caagcagaagacggcatacgaGATCGGTCTCGGCATTCCTGCTGAACCGCTCTTCCGATCT-3’.

• The 50 µl amplification reaction consisted of 0.5 µM tagged primers, 25 µl 2x Platinum™ SuperFi™ PCR Master Mix (Thermo Fisher Scientific, Catalog number 12351010), 17 µl of RNase-free water, and 3 µl of the DNA template.

• The PCR profile included an initial step for nick translation at 72 °C for 5 min, then denaturation at 98 °C for 30 s, followed by 15 cycles of 98 °C for 10 s, 64 °C for 10 s and 72 °C for 5 s, and finally at 75 °C for 5 min. After the last cycle, the PCR products were immediately cooled down to 4 °C.

• The amplified libraries were then purified with Solid Phase Reversible Immobilization (SPRI) approach using a 1.8x ratio in order to remove leftover PCR reagents and fragments smaller than 100 bp following Vesterinen et al. (2016).

• Each PCR batch included a negative control to measure the purity of reagents and the level of cross-contamination. All the PCR controls proved to negative, that is, free of contamination.

5. SIZE SELECTION AND SEQUENCING

• The DNA concentration of each purified DNA library was measured with Qubit Fluorometer, and fragments of 250–550 bp were selected for subsequent analysis using E-gel system with SizeSelect™ Agarose gel 2% (Thermo Fisher Scientific, Catalog number G661012). Briefly, the samples were loaded in the upper wells, and run down until they reached the lower wells, and then transferred into separate tubes for further processing.

• The concentration and the size profile of the size-selected DNA libraries were measured by running them on an Agilent 2100 Bioanalyzer Instrument (Agilent Technologies, Part number G2939BA) using High-Sensitivity DNA Kit (Agilent Technologies, Part number 5067-4626).

• As the DNA concentration and the purity of the DNA libraries was sub-optimal, an ammonium acetate precipitation protocol was applied to further purify and concentrate the libraries:

1. First, 5 µl of Ammonium acetate was added to 50 µl of sample.

2. Then, 138 µl of ~96% ethanol was added to the mixture and mixed by inverting the tube.

3. The mixture was incubated at -20 °C for overnight.

4. The next day, mixture was centrifuged at 15,000x g for 30 min at +4 °C.

5. The supernatant was carefully removed, the pellet was dried and finally eluted to 7 µl or RNase-free water.

• First, the DNA concentration of precipitated samples was checked with Qubit.

• The integrity and the size profile of the concentrated DNA libraries were measured again by running them using a Bioanalyzer Highsensitivity DNA Kit.

• Finally, each library pool was sequenced on one Illumina NovaSeq6000 SP flow cell using 300 cycles (2x150 paired-end) by the Turku Centre for Biotechnology, Turku, Finland.

6. BIOINFORMATICS

• The sequencing run produced 934,066,740 reads demultiplexed to the original samples by accepting one mismatch in the index.

• On average, each sample (n=263) had an output of 3,552,000 reads.
